# Complete Chloroplast Genome Sequence and Phylogenetic Analysis of *Camellia sinensis* sp. Baihaozao

**DOI:** 10.1101/2024.08.27.609997

**Authors:** Zhiyin Chen, Youpeng Zhu, Zhiming He, Hongyu Li, Jing Huang, Yihui Gong

**Affiliations:** College of Agriculture & Biotechnology, Hunan University of Humanities, Science & Technology, Loudi 417000, China; Demonstration Base for Innovation and Entrepreneurship in the Efficient Utilization of Local Characteristic Resources in Hunan Province, Loudi 417000, China; Tea Research Institute, Hunan Academy of Agricultural Sciences, Changsha 410125, China; Taohuayuan Agricultural Development Co., Ltd., Xinhua County, Loudi 417000, China

**Keywords:** Baihaozao, chloroplast genomes, structural features, phylogeny

## Abstract

Baihaozao (*Camellia sinensis* sp. Baihaozao) is esteemed in the domain of tea plant cultivation for its early harvest period, profusion of bud trichomes, and exceptional suitability for tea processing. Nevertheless, the intricate phylogenetic relationships among species within this genus remain ambiguous, primarily due to the limited availability of genomic data. This study endeavors to comprehensively investigate the genomic resources of Baihaozao by sequencing, assembling, and annotating its entire chloroplast (cp) genome. The sequencing results indicated that the total length of the Baihaozao cp genome is 157,052 base pairs (bp), with an overall guanine-cytosine (GC) content of 37.30%. The genome exhibits a typical quadripartite structure, consisting of a large single-copy region (LSC) of 86,586 bp, a small single-copy region (SSC) of 18,227 bp, and a pair of inverted repeats (IRs) totaling 52,162 bp. A total of 133 genes were identified within this genome, including 8 ribosomal RNA (rRNA) genes, 37 transfer RNA (tRNA) genes, 87 protein-coding genes, and 1 pseudogene. Furthermore, the study identified 157 simple sequence repeats (SSRs) and 90 long repeat sequences. Analysis of codon usage bias indicated that codons encoding leucine (Leu) were the most frequently utilized, whereas those encoding cysteine (Cys) were the least utilized. Examination of nucleotide diversity within the Baihaozao cp genome revealed five highly variable regions with elevated Pi values (*rps19*, *rpl32*, *ndhF*, *rpl22*, *matK*), suggesting their potential utility as molecular markers. Phylogenetic analysis of 20 cp genomes of the Theaceae family indicated a close evolutionary relationship between Baihaozao and *Camellia sinensis* OL450428.1. This study not only provides valuable data support for elucidating the maternal origin of cultivated Camellia species but also holds significant scientific implications for further exploring the phylogenetic relationships and germplasm resource utilization of Camellia plants.

## Introduction

The tea plant, classified under the genus Camellia within the family Theaceae, represents a plant resource of considerable economic and cultural importance[1]. In China, the history of tea plant cultivation extends back several millennia[2]. Contemporary research on tea plants predominantly concentrates on various domains: cultivation and breeding, plant protection[3], extraction of functional components from tea leaves, and their pharmacological properties[4, 5]. Despite notable advancements in these fields, comprehensive studies on the genomic sequencing of tea plants remain relatively limited[6]. In comparison to other species within the genus *Camellia*, the body of genomic research literature is notably sparse[7]. Baihaozao (*Camellia sinensis* sp. Baihaozao), a tea plant variety of national repute, has been officially designated as “Huacha 69” (GSCT 69). This variety is distinguished by its high yield, exceptional resistance to cold and disease, and superior quality, leading to its extensive cultivation in the Jiangnan region. Given the substantial economic importance of Baihaozao within the tea industry, comprehensive research into its genomic sequence is of considerable scientific and practical value. However, existing scientific literature on the genomic sequence of Baihaozao remains sparse, underscoring the need for further investigation and systematic study. In recent years, advancements in chloroplast (cp) genome sequencing technology, coupled with reduced costs, have rendered cp genome research a crucial tool for species identification and evolutionary studies[8]. The cp, a distinctive organelle present in green plants and algae, serves as the principal site of photosynthesis and harbors an independent genetic system characterized by a closed circular double-stranded DNA molecule[9]. This genetic system, which operates based on genetic information provided by the cell nucleus, is capable of semi-autonomous replication[10]. In higher plants, the cp genome generally manifests a quadripartite structure, consisting of a large single-copy region (LSC), a small single-copy region (SSC), and two inverted repeat sequences (IRa and IRb) [11, 12]. Despite the high conservation of sequence and structure in the cp genome across different species, its size typically varies between 115 and 165 kilobases[13, 14]. The gene content and order exhibit relative stability[15]. The structural attributes of the cp genome, especially the sequence variations in non-coding regions, offer high-resolution genetic markers for phylogenetic analysis of plant populations[16]. These markers include intergenic spacer regions, simple sequence repeats (SSRs), and single nucleotide polymorphisms[17]. Markers not only facilitate the tracing of plant origins, migrations, and evolutionary histories but are also extensively employed in the taxonomic identification of various species within a genus or distinct individuals within a species[18]. Furthermore, the uniparental inheritance and differential gene evolution rates of the cp genome provide unique advantages in plant phylogenetics and population genetics studies[19]. Consequently, research on the cp genome holds broad application prospects and significant scientific value in the field of botany.

This study utilizes Illumina high-throughput sequencing technology to perform an exhaustive sequencing and analysis of the cp genome of *Camellia sinensis* sp. Baihaozao, a species within the Theaceae family, with the objective of elucidating its genomic characteristics. The specific aims of the research are as follows: (1) Molecular Structure Analysis: Undertake a comprehensive examination of the molecular structure of the Baihaozao cp genome, encompassing its gene composition and organizational patterns; (2) Analysis of Repeat Sequences and SSR Variability: Conduct a systematic detection of repeat sequences and SSR variations within the Baihaozao cp genome to elucidate molecular-level insights into genome diversity and stability; (3) Identification of Divergence Hotspots: Investigate divergence hotspot regions within the cp genome to identify potential molecular markers, thereby establishing a foundation for future phylogenetic studies; (4) Phylogenetic Relationship Construction: Based on the complete cp genome sequence of Baihaozao, specifically the LSC, SSC, and IR, construct a phylogenetic tree of Theaceae species and analyze their phylogenetic relationships. By achieving these research objectives, this study is expected to provide valuable reference data for the identification of Theaceae species, exploration of evolutionary mechanisms, and research in the fields of population genetics and phylogenetics.

## 1. Materials and Methods

### 1.1 DNA Extraction and Quality Control of Data

Fresh young leaves of *Camellia sinensis* (L.) O. Kuntze sp. Baihaozao were collected from Taohuayuan Agricultural Development Co., Ltd., located in Xinhua County, Hunan Province, China (27°47′29.01″N, 110°51′50.86″E). These tender leaves were utilized for the extraction of the total genome (Fig. 1). The genomic DNA [20]of Baihaozao was isolated using a plant genomic DNA extraction kit (TIANGEN, DP305). The purity and concentration of the extracted DNA were evaluated through 1.0% agarose gel electrophoresis. Following verification of the DNA quality, the genomic DNA was fragmented using ultrasonic techniques. The fragmented DNA underwent a series of preparatory steps including fragment purification, end repair, 3’ end A-tailing, and adapter ligation, followed by size selection via agarose gel electrophoresis. The selected fragments were then Polymerase chain reaction (PCR) amplified to construct a sequencing library. This library was subjected to quality control assessments, and those that met the quality criteria were sequenced on an Illumina NovaSeq 6000 platform utilizing paired-end (PE150) [21]sequencing. The resulting raw data were processed using fastp v0.23.4[22] (https://github.com/OpenGene/fastp) to generate high-quality clean data. All of the aforementioned procedures were carried out by Genepioneer Biotechnologies Co. Ltd. (Nanjing, China).

**Fig. 1.**
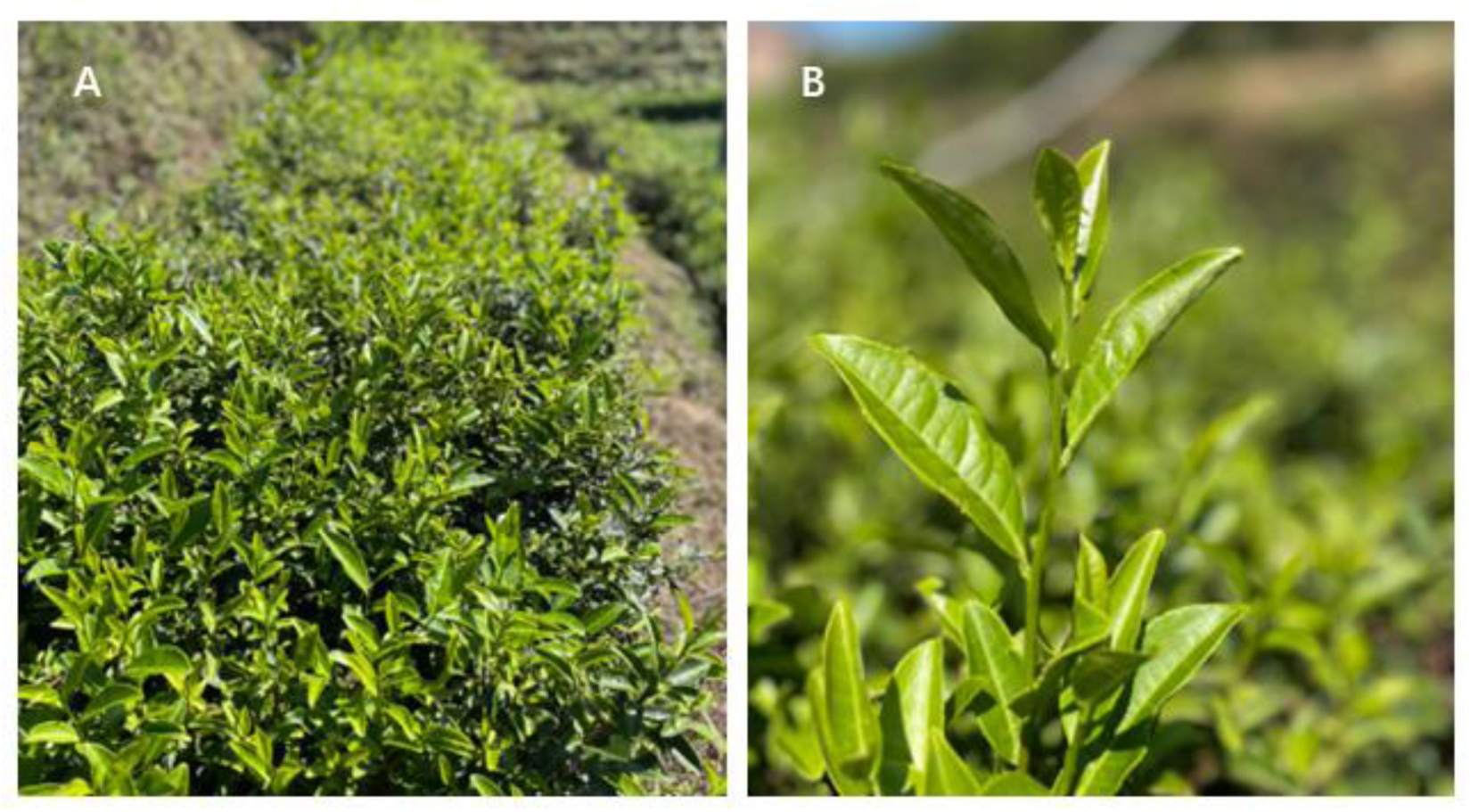
The planted field (A) and single tea plant of Baihaozao (B)

### 1.2 Genome Assembly

The sequencing reads were aligned to a custom-built cp genome database using Bowtie2 v2.2.4 (http://bowtie-bio.sourceforge.net/bowtie2/index.shtml) in the very-sensitive-local mode. The aligned sequences were subsequently designated as the cp genome sequencing sequences (cpDNA sequences) for the project samples. The core assembly module employed SPA des v3.10.1[23, 24] (http://cab.spbu.ru/software/spades/) for the assembly of the cp genome. Quality control of the assembled genomes was conducted utilizing the reference sequence *Camellia sinensis* (NCBI accession number OL450397.1.gbk), which is accessible via the NCBI database at https://www.ncbi.nlm.nih.gov/nuccore/. It is important to note that different samples may require the use of distinct reference sequences.

### 1.3 Structure of Cp Genes and Construction of Genetic Maps

Initially, cp coding sequences (CDS) were annotated utilizing Prodigal v2.6.3[25] (https://www.github.com/hyattpd/Prodigal). Concurrently, rRNA predictions were performed using HMMER v3.1b2 (http://www.hmmer.org/), and tRNA predictions were executed with ARAGORN v1.2.38 [26](http://www.ansikte.se/ARAGORN/). Subsequently, the assembled sequences were compared against gene sequences of closely related species available on NCBI, employing BLAST v2.6 (https://blast.ncbi.nlm.nih.gov/Blast.cgi) to derive a secondary set of annotation results. Next, discrepancies between the two annotation results were manually scrutinized to eliminate erroneous and redundant annotations, thus delineating the boundaries of multiple exons and finalizing the annotation. The cp genome map was generated utilizing OGDRAW[27] (https://chlorobox.mpimp-golm.mpg.de/OGDraw.html). Afterward, discrepancies between the two annotation results were manually examined to remove erroneous and redundant annotations, thereby determining the boundaries of multiple exons and achieving the final annotation. The cp genome map was generated using OGDRAW (https://chlorobox.mpimp-golm.mpg.de/OGDraw.html).

### 1.4 Analysis of Genomic Structural Annotation

The method for calculating Relative Synonymous Codon Usage (RSCU) is defined as follows: (the frequency of a specific codon encoding a particular amino acid / the total frequency of all codons encoding that amino acid) / (1 / the number of synonymous codons for that amino acid). This metric quantifies the observed frequency of a codon relative to its expected frequency under uniform codon usage. A custom Perl script was employed to filter unique coding sequences (CDS), selecting one representative from multiple copies, and to perform subsequent calculations. Repetitive sequences were identified using vmatch v2.3.0 (http://www.vmatch.de/) in conjunction with additional Perl scripts. The parameters for this identification included a minimum length of 30 base pairs and a Hamming distance of 3, with four identification forms: forward, palindromic, reverse, and complement. cpSSRs analysis was conducted using MISA v1.0 (MIcroSAtellite identification tool, https://webblast.ipk-gatersleben.de/misa/).

### 1.5 Analysis of Nucleotide Diversity and Nucleotide Substitution Rate (Ka/Ks)

Gene sequences were aligned using MAFFT v7.427[28] (https://mafft.cbrc.jp/alignme nt/software), and the Ka/Ks values of the genes were calculated using Ka/Ks Calculator v2.0 [29] (https://sourceforge.net/projects/kakscalculator2/).

### 1.6 Analysis of Nucleotide Diversity(π)

A global alignment of homologous gene sequences from different species was conducted using the MAFFT software in --auto mode. The π (pi) values for each gene were subsequently calculated using DNAsp5 [30](http://www.ub.edu/dnasp/).

### 1.7 Analysis of Comparative Genomics

The cp genome exhibits a circular configuration, characterized by four demarcations between the IR and LSC and SSC regions, specifically designated as LSC-IRb, IRb-SSC, SSC-IRa, and IRa-LSC. Throughout the evolutionary trajectory of the genome, the boundaries of the IR regions are subject to expansion and contraction, facilitating the translocation of certain genes into either the IR or single-copy regions. The delineation of these boundaries was graphically represented utilizing the CPJSdraw tool available on the Gene Pioneer cloud platform (http://cloud.genepioneer.com:9929/#/tool/alltool/detail/296).

For the purpose of phylogenetic analysis, a whole-genome approach was utilized, standardizing the circular sequences to a common starting point. Interspecies sequences were aligned using MAFFT version 7.427 in --auto mode. The aligned sequences were subsequently analyzed with RAxML version 8.2.10 [31](https://cme.h-its.org/exelixis/software.html), employing the GTRGAMMA model and rapid Bootstrap analysis (bootstrap = 1000) to construct a maximum likelihood phylogenetic tree. Comparative genomic analysis was conducted using Mauve (https://darlinglab.org/mauve/mauve.html) with default settings.

## 2. Results

### 2.1 Genome Structure of the Baihaozao cp Genome

The cp genome of Baihaozao spans 157,052 base pairs (bp) and exhibits a typical circular quadripartite structure. This structure comprises an 18,277 bp SSC region, an 86,586 bp LSC region, and two IR regions, designated IRa and IRb, which collectively total 52,162 bp (Fig. 2 and Table 1). The overall guanine-cytosine (GC) content of the cp genome is 37.30%, with a non-uniform distribution across different regions. Specifically, the GC content is higher in the IR regions (42.95%) compared to the LSC (35.33%) and SSC (30.55%) regions. Furthermore, the cp genome of Baihaozao contains 133 predicted functional genes, including 87 protein-coding genes, 37 tRNA genes, 8 rRNA genes, and 1 pseudogene.

**Fig. 2.**
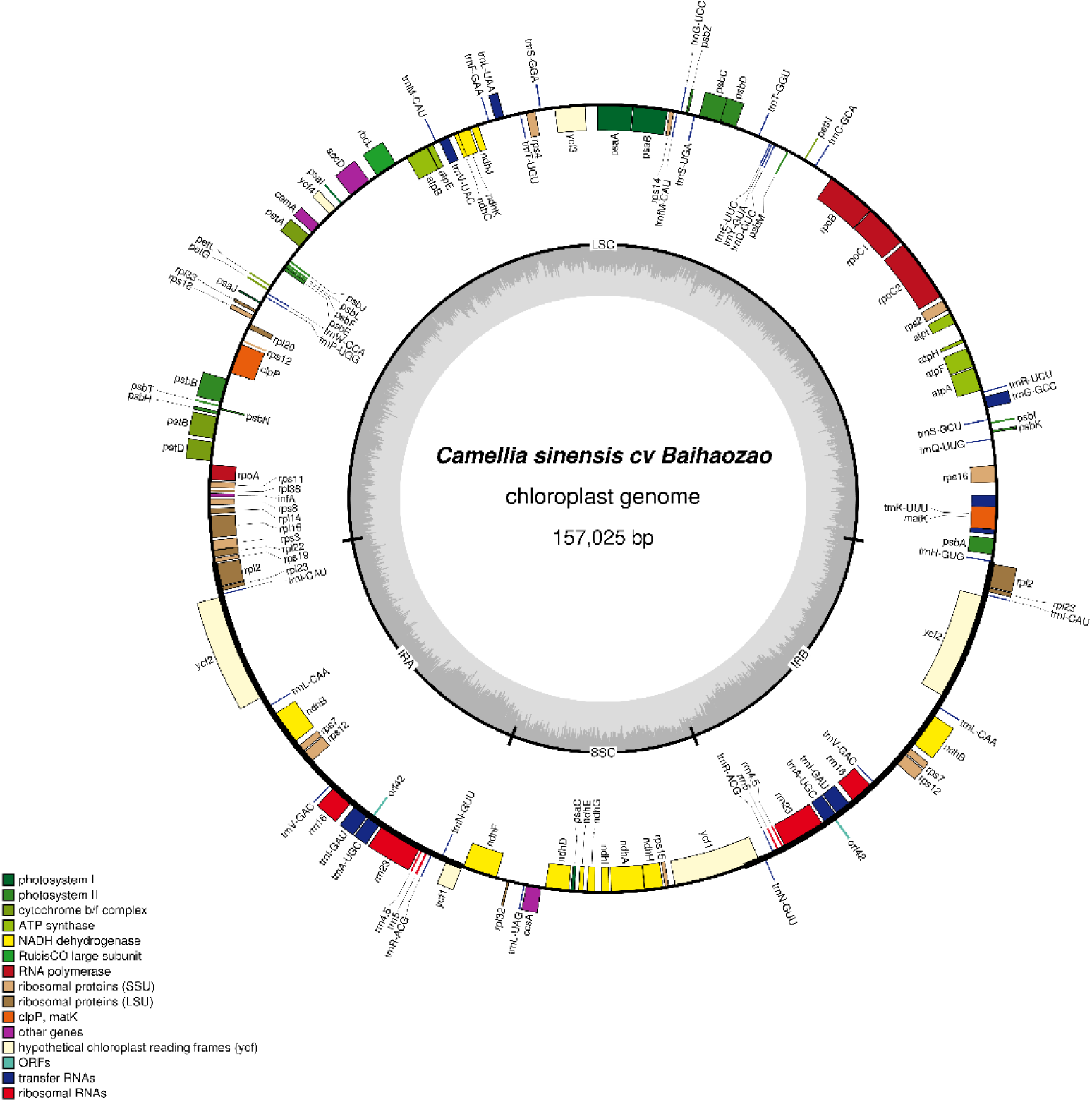
Gene map for cp genome of the Baihaozao.

**Table 1.** Characteristics of the cp genome of Baihaozao.

| Category | Item | Describe |
| --- | --- | --- |
| Chloroplast genome structure | Cp gene/bp | 157,025 |
|  | LSC/bp | 86,586 |
|  | SSC/bp | 18,277 |
|  | IRa/IRb/bp | 52,162 |
| Gene composition | Cp gene | 133 |
|  | tRNA | 37 |
|  | rRNA | 8 |
|  | mRNA | 87 |
|  | pseudo | 1 |
| GC Content (%) | Cp gene | 37.30 |
|  | LSC | 35.33 |
|  | SSC | 30.55 |
|  | IRa/IRb | 42.95 |

As illustrated in Table 2, the cp genome’s IR regions encompass 19 duplicated genes, which include 8 protein-coding genes (*ndhB*, *rpl2*, *rpl23*, *rps12*, *rps7*, *orf42*, *ycf1*, and *ycf2*), 7 tRNA genes (*trnA-UGC*, *trnI-CAU*, *trnI-GAU*, *trnL-CAA*, *trnN-GUU*, *trnR-ACG*, and *trnV-GAC*), and 4 rRNA genes (*rrn16*, *rrn23*, *rrn4.5*, and *rrn5*). Furthermore, among the 133 genes, 14 genes (*ndhA*, *ndhB*, *petB*, *petD*, *rpl16*, *rpl2*, *rps16*, *rpoC1*, *trnA-UGC*, *trnG-GCC*, *trnI-GAU*, *trnK-UUU*, *trnL-UAA*, and *trnV-UAC*) contain a single intron, 3 genes (*rps12*, *clpP*, and *ycf3*) contain two introns, and there is one pseudogene (*ycf1*).

**Table 2.**
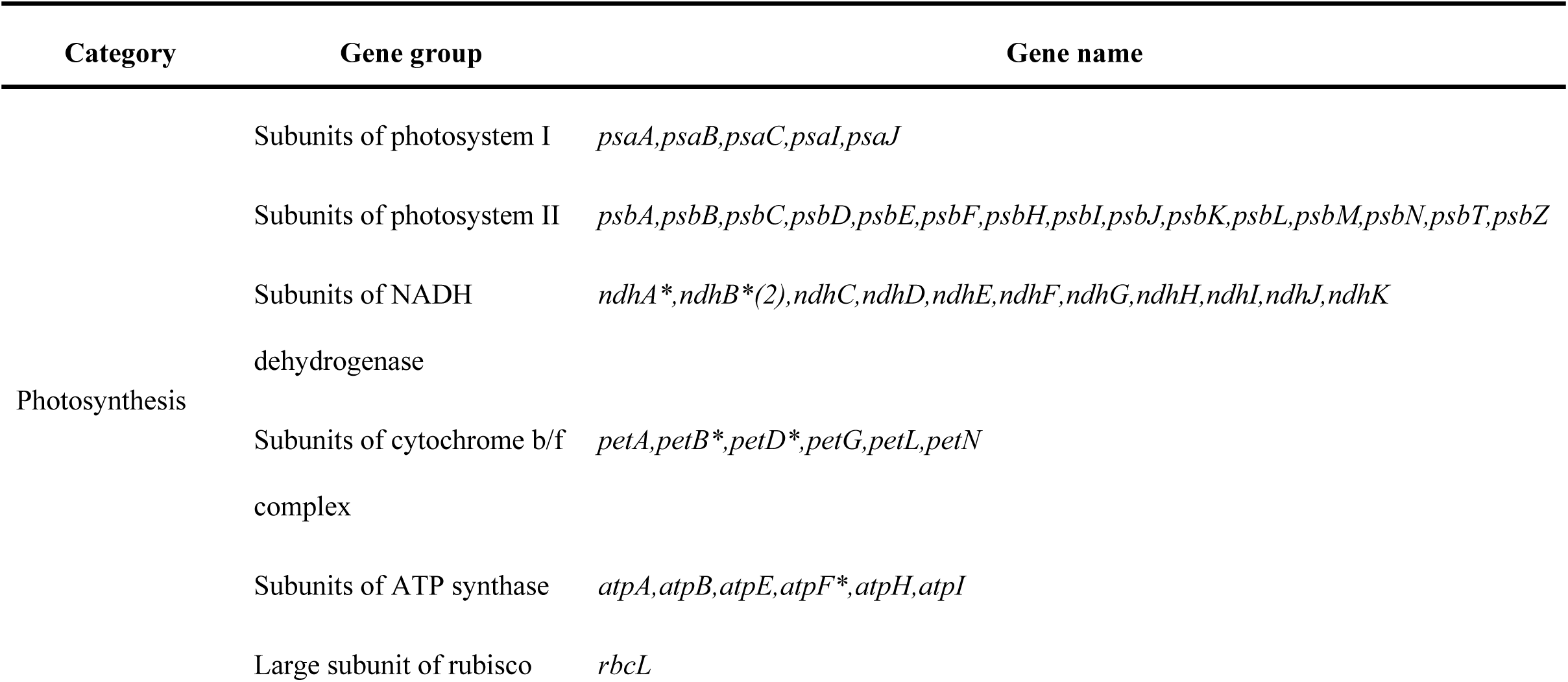
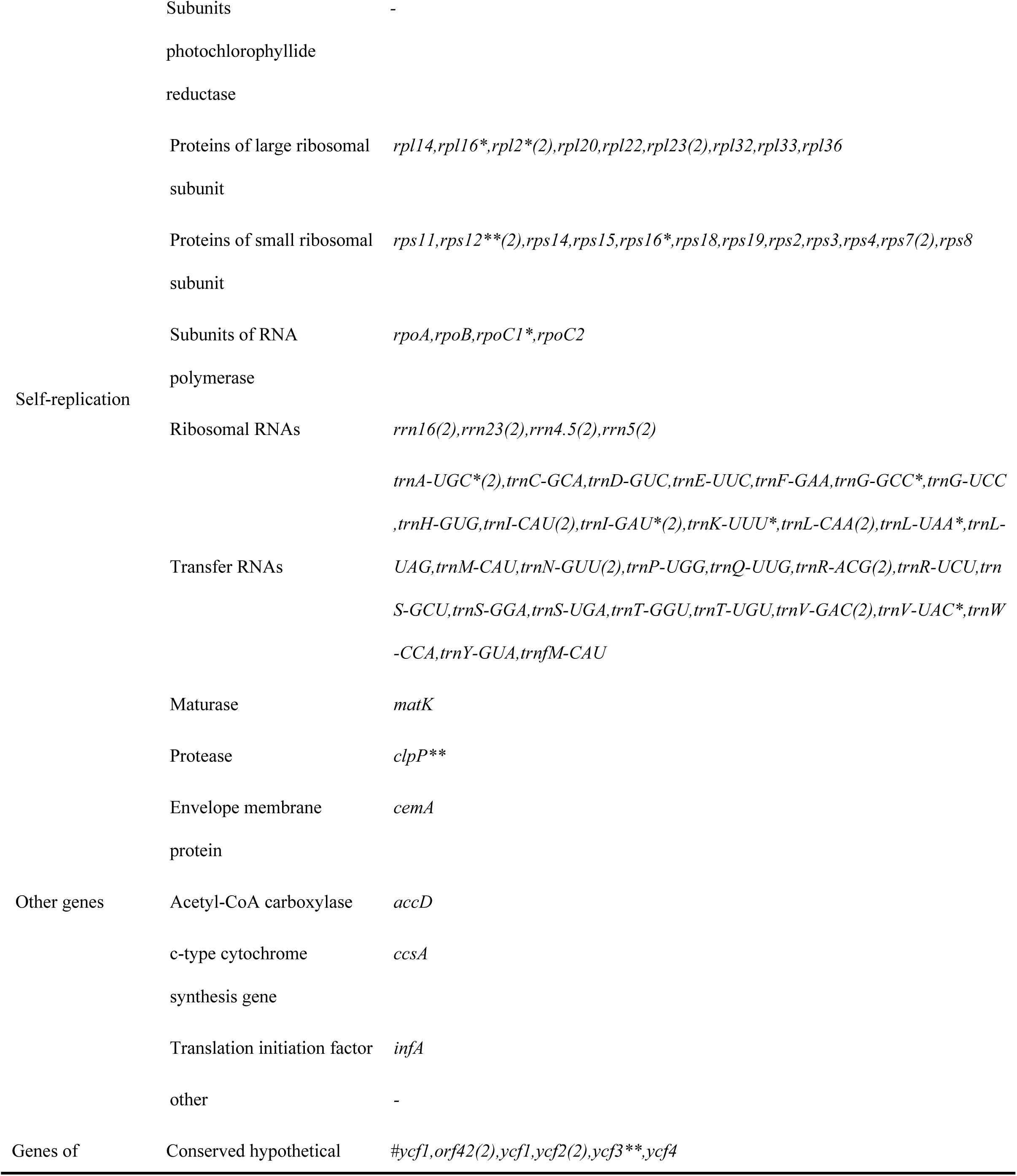
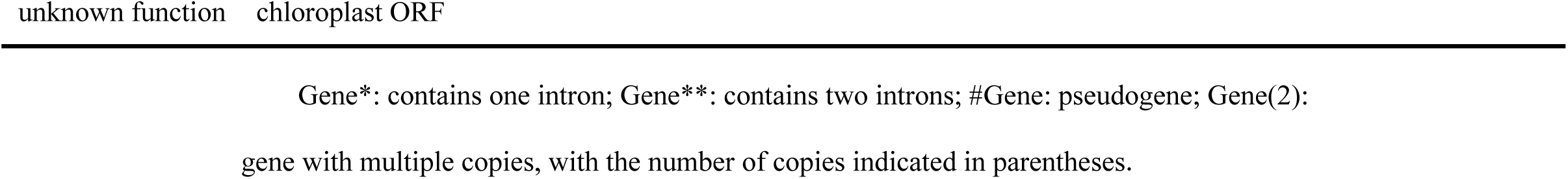
Genes present in cp genome of ‘Baihaozao’.

### 2.2 Codon Usage Bias

RSCU refers to the unequal usage of synonymous codons, a phenomenon influenced by natural selection, species-specific mutations, and genetic drift. Analysis of the Baihaozao genome revealed that protein-coding genes utilize a total of 26,471 codons, including 3 stop codons (Table 3 and Fig. 3). Among these, the codon for Leucine (Leu) is the most prevalent, accounting for 2,740 codons (10.35%), followed by Isoleucine (Ile) with 2,284 codons (8.63%), and Serine (Ser) with 2,068 codons (7.81%). Conversely, Cysteine (Cys) is the least prevalent amino acid, encoded by a mere 296 codons, constituting 1.12% of the total. Among the codons, 32 (72.7%) exhibit an RSCU value exceeding 1. The codon AUG, encoding Methionine (Met), is the most preferentially utilized, with an RSCU value of 6.9566, followed by UUA, which encodes Leucine, with an RSCU value of 1.8984.

**Fig. 3.**
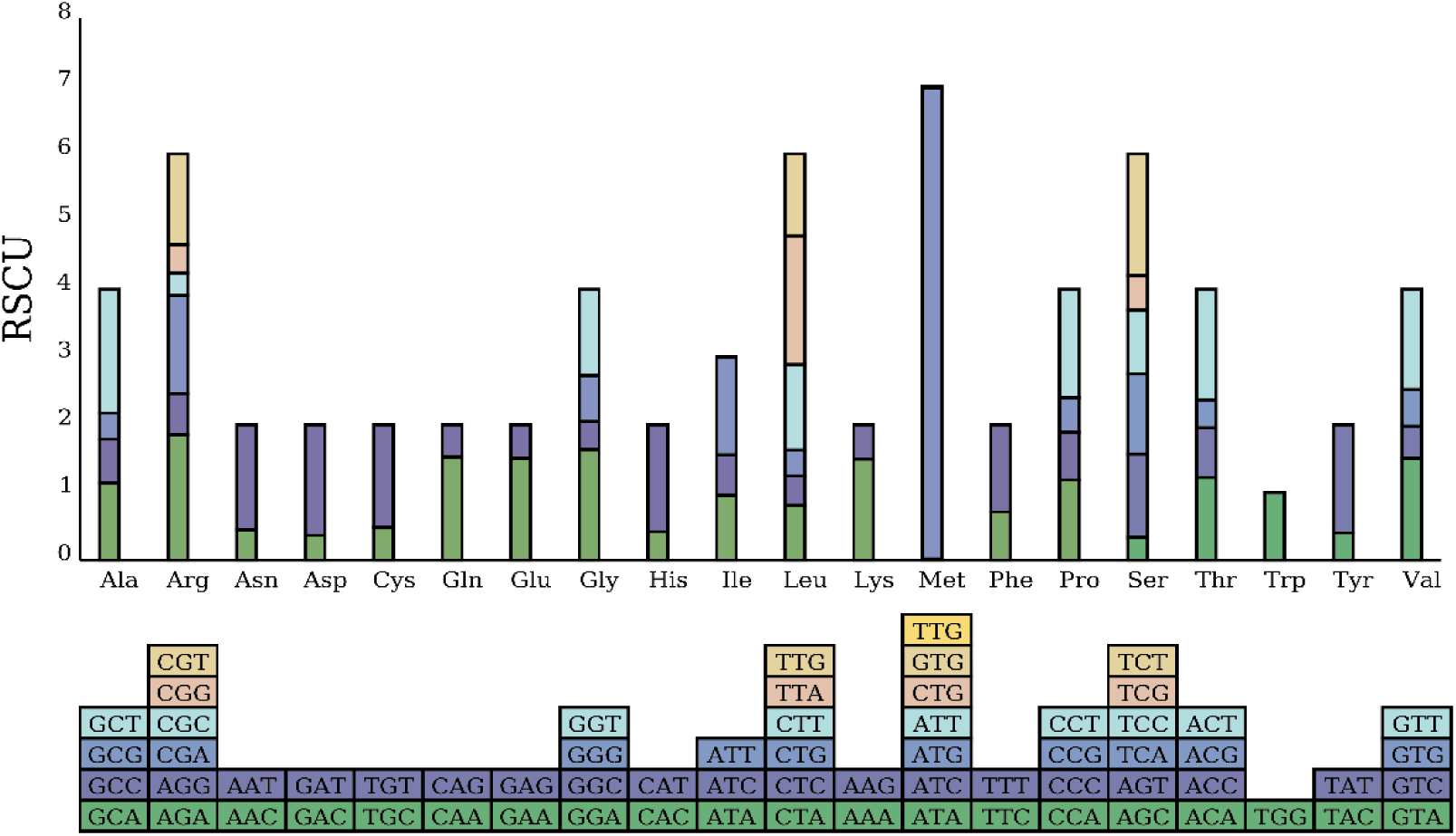
RSCU bar chart. Note: The boxes below represent all codons encoding each amino acid, while the height of the columns above indicates the total RSCU values for all codons.

**Table 3.**
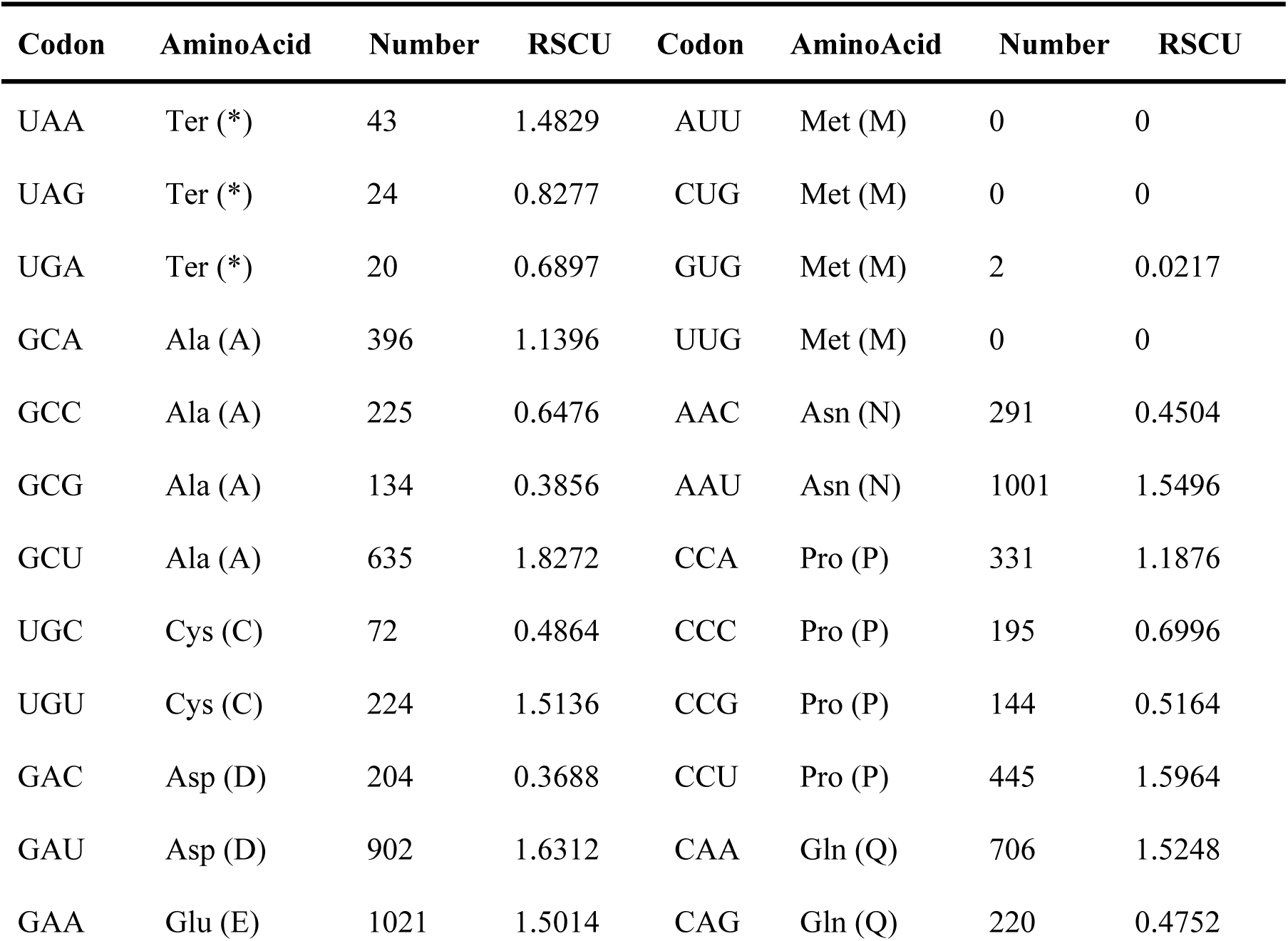
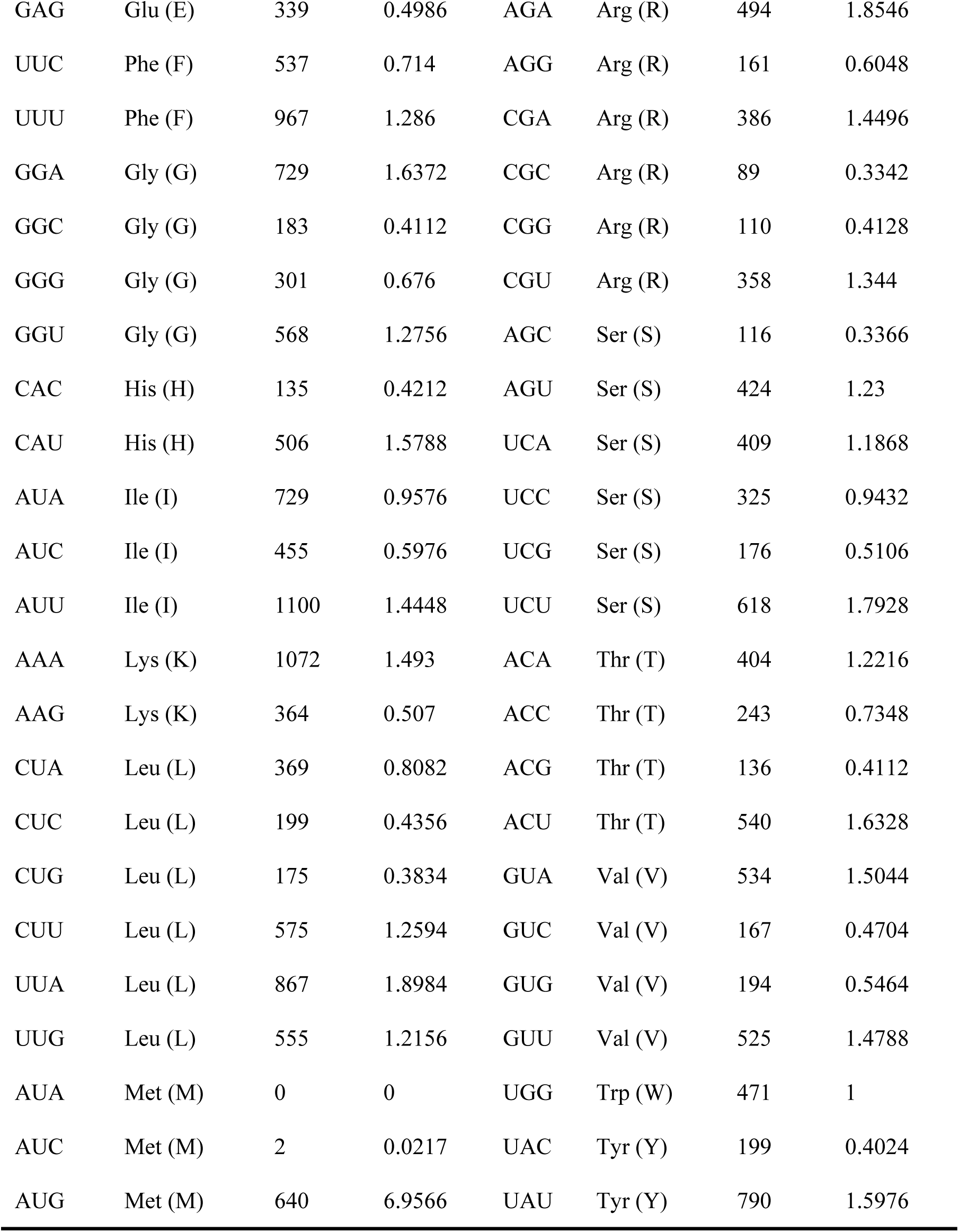
Codon list of Baihaozao.

| Codon | AminoAcid | Number | RSCU | Codon | AminoAcid | Number | RSCU |
| --- | --- | --- | --- | --- | --- | --- | --- |
| UAA | Ter (*) | 43 | 1.4829 | AUU | Met (M) | 0 | 0 |
| UAG | Ter (*) | 24 | 0.8277 | CUG | Met (M) | 0 | 0 |
| UGA | Ter (*) | 20 | 0.6897 | GUG | Met (M) | 2 | 0.0217 |
| GCA | Ala (A) | 396 | 1.1396 | UUG | Met (M) | 0 | 0 |
| GCC | Ala (A) | 225 | 0.6476 | AAC | Asn (N) | 291 | 0.4504 |
| GCG | Ala (A) | 134 | 0.3856 | AAU | Asn (N) | 1001 | 1.5496 |
| GCU | Ala (A) | 635 | 1.8272 | CCA | Pro (P) | 331 | 1.1876 |
| UGC | Cys (C) | 72 | 0.4864 | CCC | Pro (P) | 195 | 0.6996 |
| UGU | Cys (C) | 224 | 1.5136 | CCG | Pro (P) | 144 | 0.5164 |
| GAC | Asp (D) | 204 | 0.3688 | CCU | Pro (P) | 445 | 1.5964 |
| GAU | Asp (D) | 902 | 1.6312 | CAA | Gln (Q) | 706 | 1.5248 |
| GAA | Glu (E) | 1021 | 1.5014 | CAG | Gln (Q) | 220 | 0.4752 |
| GAG | Glu (E) | 339 | 0.4986 | AGA | Arg (R) | 494 | 1.8546 |
| UUC | Phe (F) | 537 | 0.714 | AGG | Arg (R) | 161 | 0.6048 |
| UUU | Phe (F) | 967 | 1.286 | CGA | Arg (R) | 386 | 1.4496 |
| GGA | Gly (G) | 729 | 1.6372 | CGC | Arg (R) | 89 | 0.3342 |
| GGC | Gly (G) | 183 | 0.4112 | CGG | Arg (R) | 110 | 0.4128 |
| GGG | Gly (G) | 301 | 0.676 | CGU | Arg (R) | 358 | 1.344 |
| GGU | Gly (G) | 568 | 1.2756 | AGC | Ser (S) | 116 | 0.3366 |
| CAC | His (H) | 135 | 0.4212 | AGU | Ser (S) | 424 | 1.23 |
| CAU | His (H) | 506 | 1.5788 | UCA | Ser (S) | 409 | 1.1868 |
| AUA | Ile (I) | 729 | 0.9576 | UCC | Ser (S) | 325 | 0.9432 |
| AUC | Ile (I) | 455 | 0.5976 | UCG | Ser (S) | 176 | 0.5106 |
| AUU | Ile (I) | 1100 | 1.4448 | UCU | Ser (S) | 618 | 1.7928 |
| AAA | Lys (K) | 1072 | 1.493 | ACA | Thr (T) | 404 | 1.2216 |
| AAG | Lys (K) | 364 | 0.507 | ACC | Thr (T) | 243 | 0.7348 |
| CUA | Leu (L) | 369 | 0.8082 | ACG | Thr (T) | 136 | 0.4112 |
| CUC | Leu (L) | 199 | 0.4356 | ACU | Thr (T) | 540 | 1.6328 |
| CUG | Leu (L) | 175 | 0.3834 | GUA | Val (V) | 534 | 1.5044 |
| CUU | Leu (L) | 575 | 1.2594 | GUC | Val (V) | 167 | 0.4704 |
| UUA | Leu (L) | 867 | 1.8984 | GUG | Val (V) | 194 | 0.5464 |
| UUG | Leu (L) | 555 | 1.2156 | GUU | Val (V) | 525 | 1.4788 |
| AUA | Met (M) | 0 | 0 | UGG | Trp (W) | 471 | 1 |
| AUC | Met (M) | 2 | 0.0217 | UAC | Tyr (Y) | 199 | 0.4024 |
| AUG | Met (M) | 640 | 6.9566 | UAU | Tyr (Y) | 790 | 1.5976 |

### 2.3 SSRs and Interspersed Repeats

Repeat sequences are DNA sequences that are reiterated hundreds or thousands of times at various loci within the genome. These repeat sequences can be broadly categorized into two primary types: tandem repeat sequences and interspersed repeat sequences. Tandem repeat sequences, also known as SSRs, typically consist of short motifs ranging from one to six nucleotides in length, repeated in a head-to-tail manner up to tens of nucleotides long. In contrast, interspersed repeat sequences, commonly referred to as transposons, are dispersed throughout the genome. In the context of the cp genome, SSR markers are designated as cpSSR markers. Our analysis of the Baihaozao cp genome identified a total of 247 SSRs. This includes 157 single nucleotide repeats, 4 dinucleotide repeats, and 72 other types of repeats. Seventy of these SSRs were equal to or exceeded 10 bp in length. The longest SSR identified was a polynucleotide repeat (CTTTTTT and AAAAAG) with a length of 18 bp. A total of 233 SSRs were A/T-based, constituting 64.5% of all SSRs (Fig. 4A). Additionally, the LSC region contained 142 SSRs (57.5%), the IR region contained 50 SSRs (20.2%), and the SSC region contained 55 SSRs (22.3%) (Fig. 4B). Further analysis revealed that 94 SSRs were located in exonic regions (38.06%), 35 SSRs were situated in intronic regions (14.17%), and 118 SSRs were found in intergenic regions (47.78%) (Fig. 4C).

**Fig. 4.**
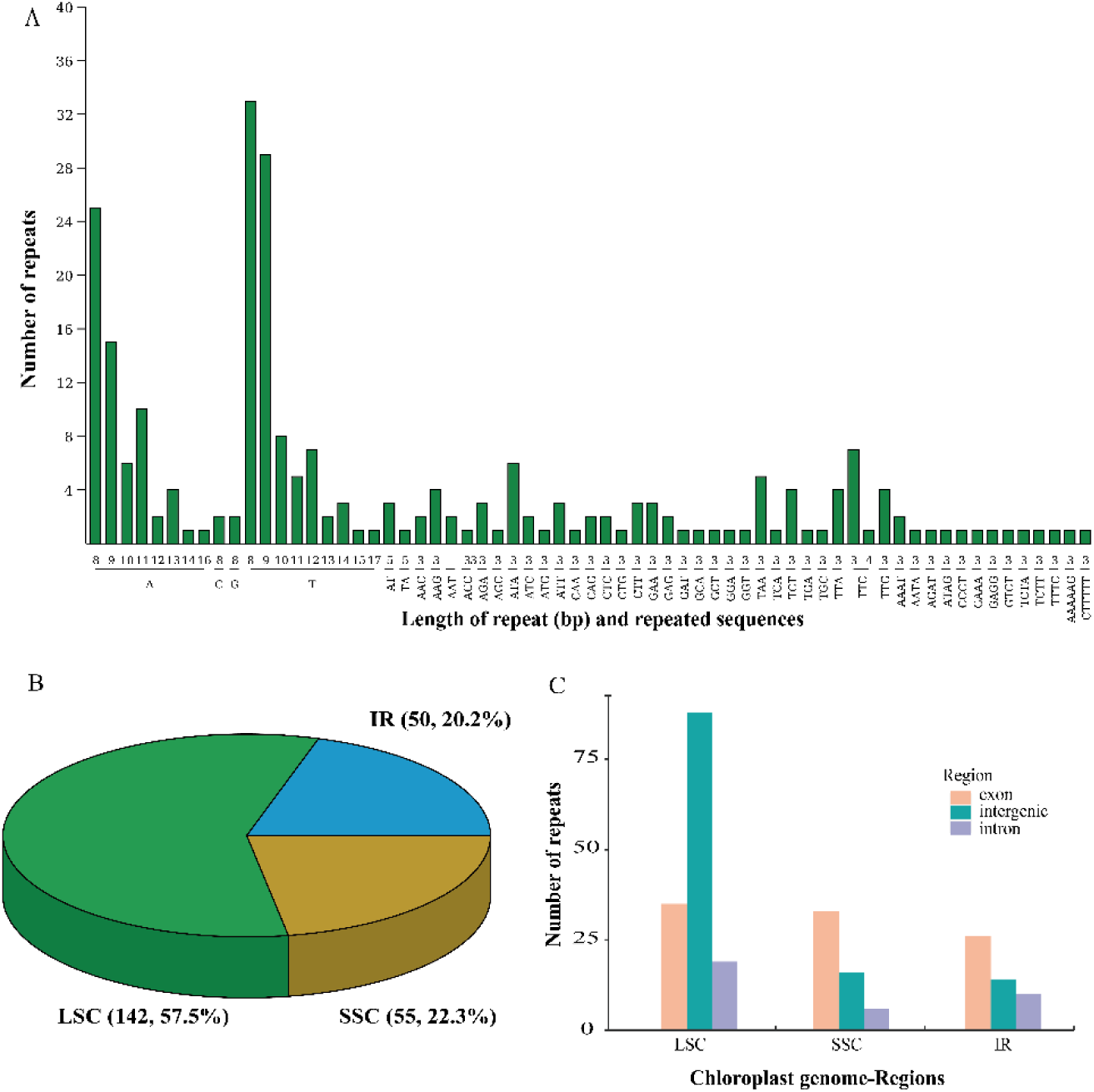
SSR Analysis of Baihaozao. Note: (A) SSR type: The horizontal axis represents the SSR repeat units, while the vertical axis indicates the number of repeat units. (B) The quantity of SSRs across various regions. (C) The distribution of SSRs within gene regions.

Scattered repeat sequences represent a distinct category of repetitive elements, differing from tandem repeat sequences by their dispersed distribution throughout the genome. In the Baihaozao cp genome, there are 40 identified scattered repeat sequences, comprising primarily 20 forward repeats and 20 palindromic repeats, with no occurrences of reverse or complementary repeats. Within the forward repeats, sequences of 42 bp were the most prevalent, accounting for five instances (25%). Similarly, among the palindromic repeats, sequences of 30 bp were the most frequent, with six instances (30%). Among the various scattered repeats, the 30 bp sequence was the most prevalent, accounting for 9 instances (22.5%). This was followed by repeats of 42 bp and 60 bp, each observed 7 times (17.5%) (refer to Fig. 5).

**Fig. 5.**
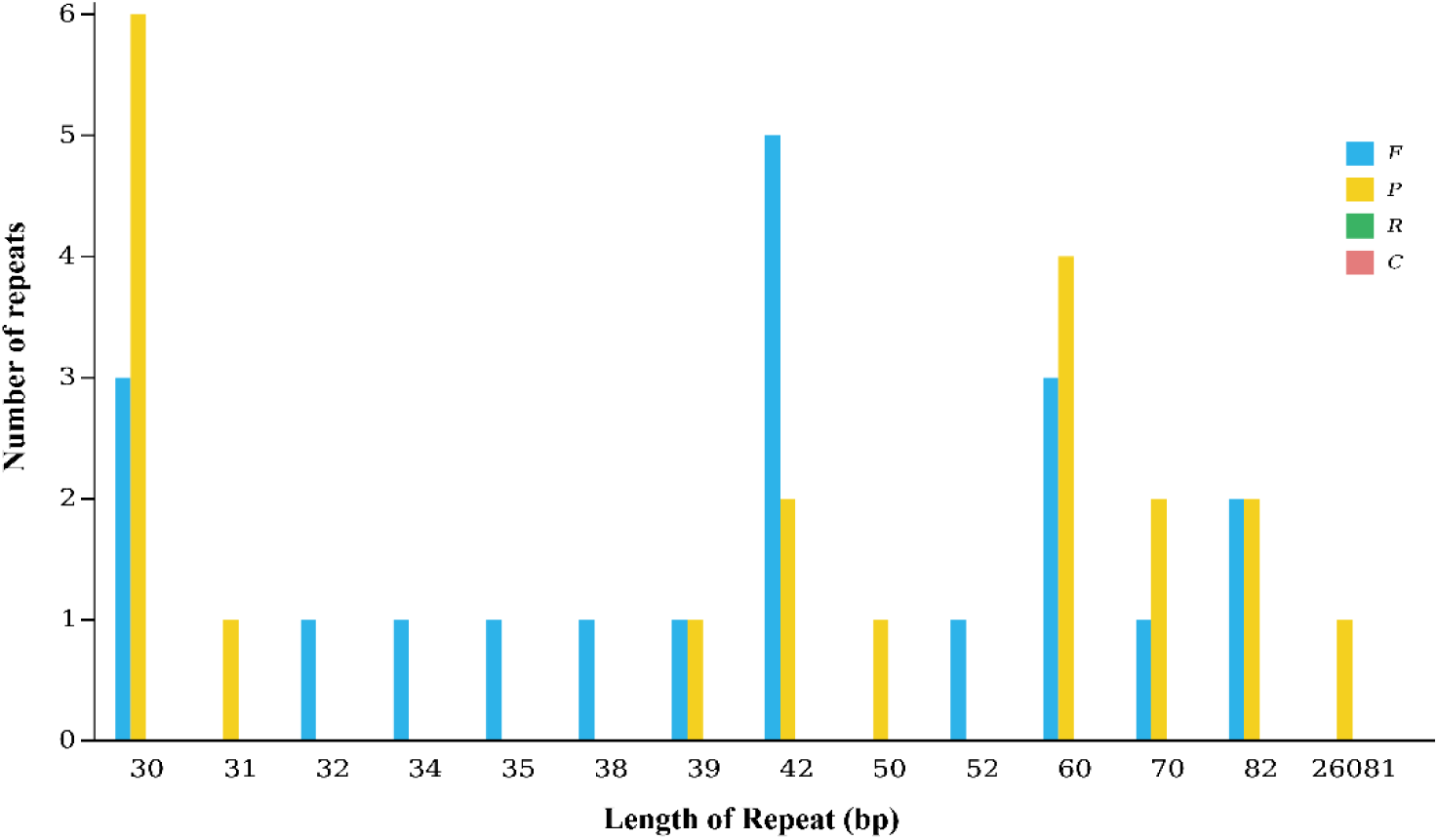
Scattered Repeat Sequence Statistics. Note: The horizontal axis represents the length of the scattered repeat sequences, while the vertical axis denotes the number of scattered repeat sequences. The abbreviations used are as follows: F for forward repeats, P for palindromic repeats, R for reverse repeats, and C for complementary repeats.

### 2.4 Ka/Ks Analysis

To elucidate gene characterization, the Ka/Ks substitution ratios of protein-coding genes were calculated across seven Camellia species. Using Baihaozao as a reference, the Ka and Ks variations in Baihaozao’s cp genome were compared with those of six other Camellia species (Fig. 6). This comparison identified a total of 197 protein-coding genes within these seven cp genomes suitable for diversity analysis. The majority of these coding genes exhibited Ka/Ks ratios lower than 1 or could not be determined due to zero Ka or Ks values, indicating their conserved nature. The *rbcL* gene exhibited the highest Ka/Ks ratio (2.59013) in MW801387, indicating rapid evolutionary changes. Additionally, five highly variable regions were identified. The *cemA* gene demonstrated Ka/Ks ratios of 1.16848 and 1.94357 in MH753078 and NC_041471, respectively. Furthermore, the *ndhD*, *psbH*, *rpl22*, and *rpoC2* genes showed Ka/Ks values exceeding 1 in MW801387, OL449838, OL449838, and OQ556867, respectively, with specific values of 1.51867, 1.0239, 1.06399, and 1.85814. These elevated Ka/Ks ratios suggest that these genes are under positive selection, whereas the remaining genes appear to be subject to negative selection.

**Fig. 6.**
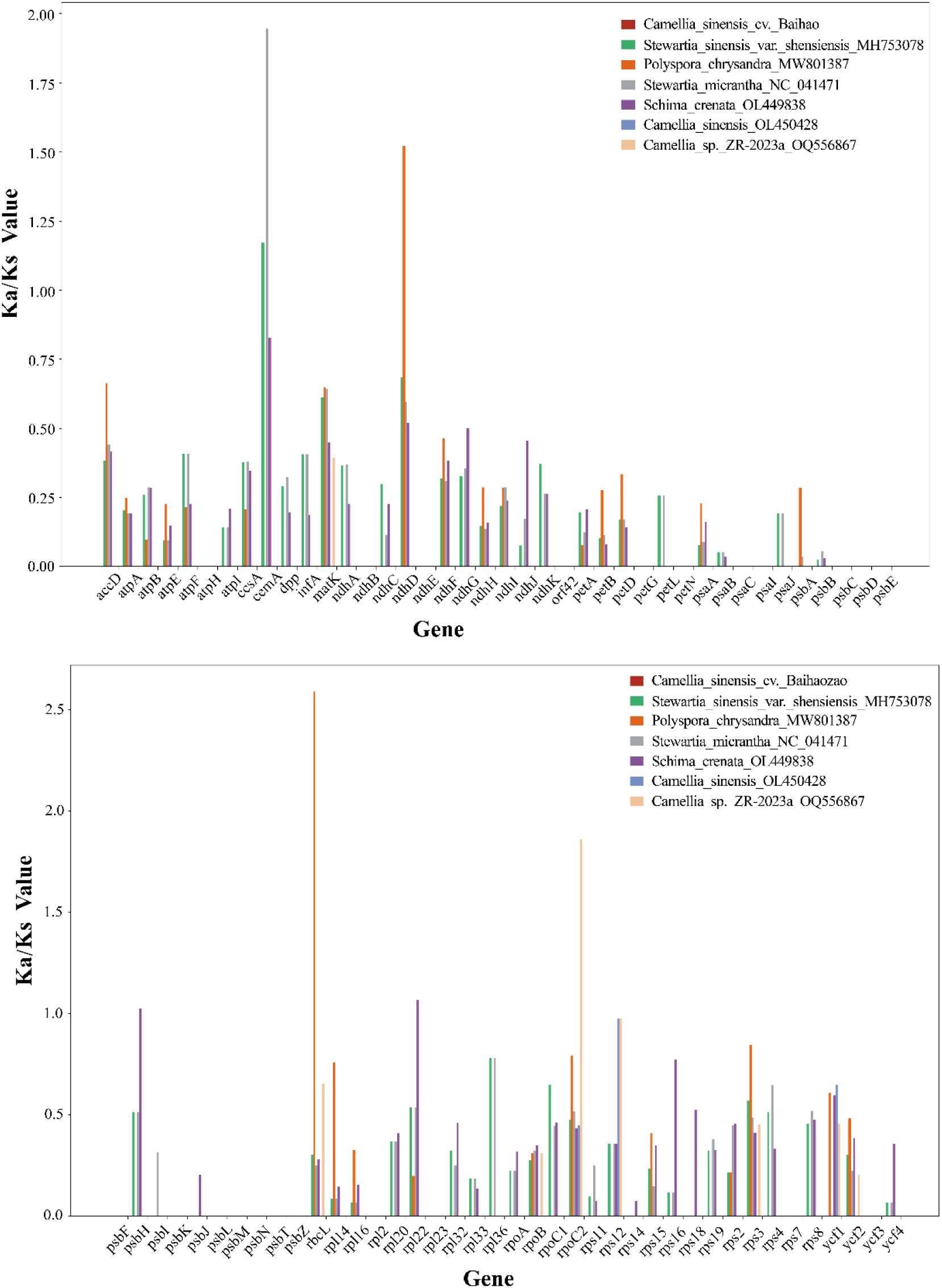
The 197 protein-coding genes in the cp genome of Baihaozao and 7 other Camellia species were analyzed for ka/ks ratios.

### 2.5 Analysis of Nucleic Acid Diversity (π)

The analysis of nucleic acid diversity, denoted as π, quantifies the extent of variation in nucleic acid sequences across different species. Regions exhibiting higher levels of variation may serve as valuable molecular markers for studies in population genetics. We analyzed the Pi values across 114 loci within the cp genome. The Pi values for the cp genome sequences ranged from 0 to 0.02675, with a mean value of 0.00713 (Fig. 7 and Supplementary Table S1). Specifically, the mean Pi value for the SSC region was 0.01261, for the LSC region was 0.00757, and for the IR region was 0.00080. These results indicate that the SSC region exhibited the highest nucleotide diversity, whereas the IR region demonstrated the lowest nucleotide diversity, suggesting a more conserved nature in the IR region. Furthermore, five genes exhibiting high Pi values were identified: *rps19* (0.02675), *rpl32* (0.02513), *ndhF* (0.02057), *rpl22* (0.01938), and *matK* (0.01891). Specifically, *rps19*, *rpl22*, and *matK* are situated in the LSC region, whereas *rpl32* and *ndhF* are located in the SSC region. These findings indicate that the *rps19*, *rpl32*, *ndhF*, *rpl22*, and *matK* loci demonstrate significant variability at the species level (Pi > 0.018) and hold potential for development as barcodes for the early identification of Baihaozao.

**Fig. 7.**
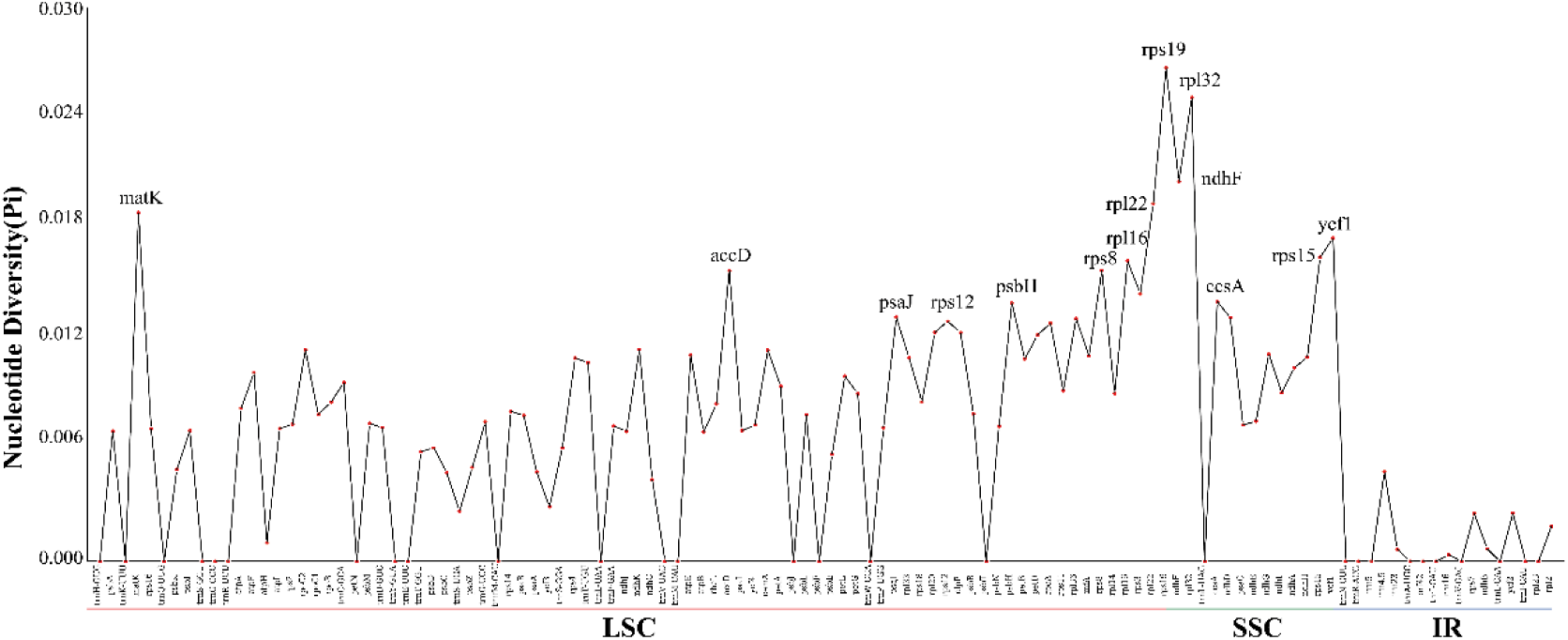
Line graph of gene pi values. Note: Horizontal coordinates indicate gene names and vertical coordinates indicate pi values.

### 2.6 Comparative Genomics

The cp genome is characterized by a circular structure, encompassing four distinct boundaries between the IR regions and the LSC and SSC regions: specifically, LSC-IRb, IRb-SSC, SSC-IRa, and IRa-LSC. Throughout the course of genomic evolution, these IR boundaries experience expansions and contractions, facilitating the translocation of certain genes into either the IR regions or the single-copy regions. Additionally, we conducted an analysis of the IR/LSC and IR/SSC boundary regions, as illustrated in Fig. 8. The location of the *rps19* gene was consistent across all seven *Camellia sinensis* species, spanning both the LSC and IRb binding regions and positioned 6-46 bp away from these regions, with the exception of *Stewartia micrantha* NC_041471, where the *rps19* gene is situated within the LSC region. Similarly, the *trnH* genes in the seven *Camellia sinensis* species are located in the LSC region and are 1-14 bp from the IRa/LSC boundary, except for *Stewartia micrantha* NC_041471, where the *trnH* gene is 30 bp from the IRa/LSC boundary. The *ndhF* genes of eight *Camellia sinensis* species are situated in the SSC region, positioned 3-66 base pairs from the IRb/SSC boundary. In contrast, the *Schima crenata* OL449838 exhibits the *ndhF* gene immediately adjacent to the IRb and SSC boundary region. Furthermore, the *trnN* genes of the eight *Camellia sinensis* species are located within the IRa and IRb regions. These findings suggest that the cp genome sequences of the eight Camellia sinensis species are conserved.

**Fig. 8.**
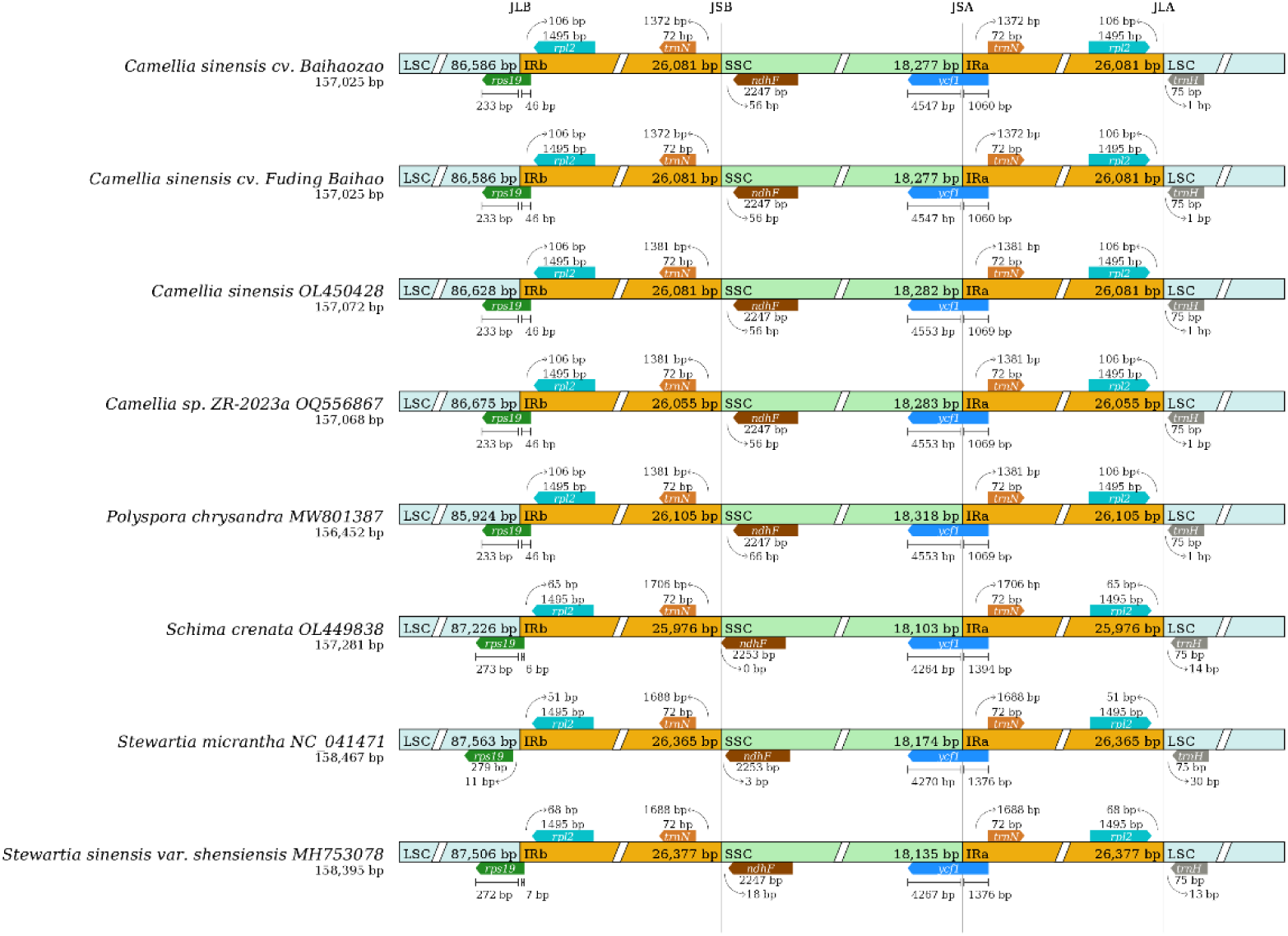
Analysis of cp IR boundary changes. Note: The thin lines represent the connection points of each region, and the figure presents information about the genes near the connection points.

The results of Mauve’s multiple comparison analysis revealed the presence of four Locally Collinear Blocks (LCBs) within the cp genomes of eight species, specifically *Polvspora chrvsandra* MW801387, *Schima crenata* QL449838, *Stewartia micrantha* NC 041471, and *Stewartia sinensis* var. shensiensis MH753078. These LCBs were identified in two genomic regions spanning 85,000-115,000 and 130,000-160,000 base pairs, respectively, indicating a high degree of genomic similarity among these species (Fig. 9). The comparative analysis revealed no rearrangements or inversions between the genomes. However, mutations were detected within the region spanning positions 5,000 to 10,000, which is characterized by a high degree of gene sequence variation in the aligned cp genomes.

**Fig. 9.**
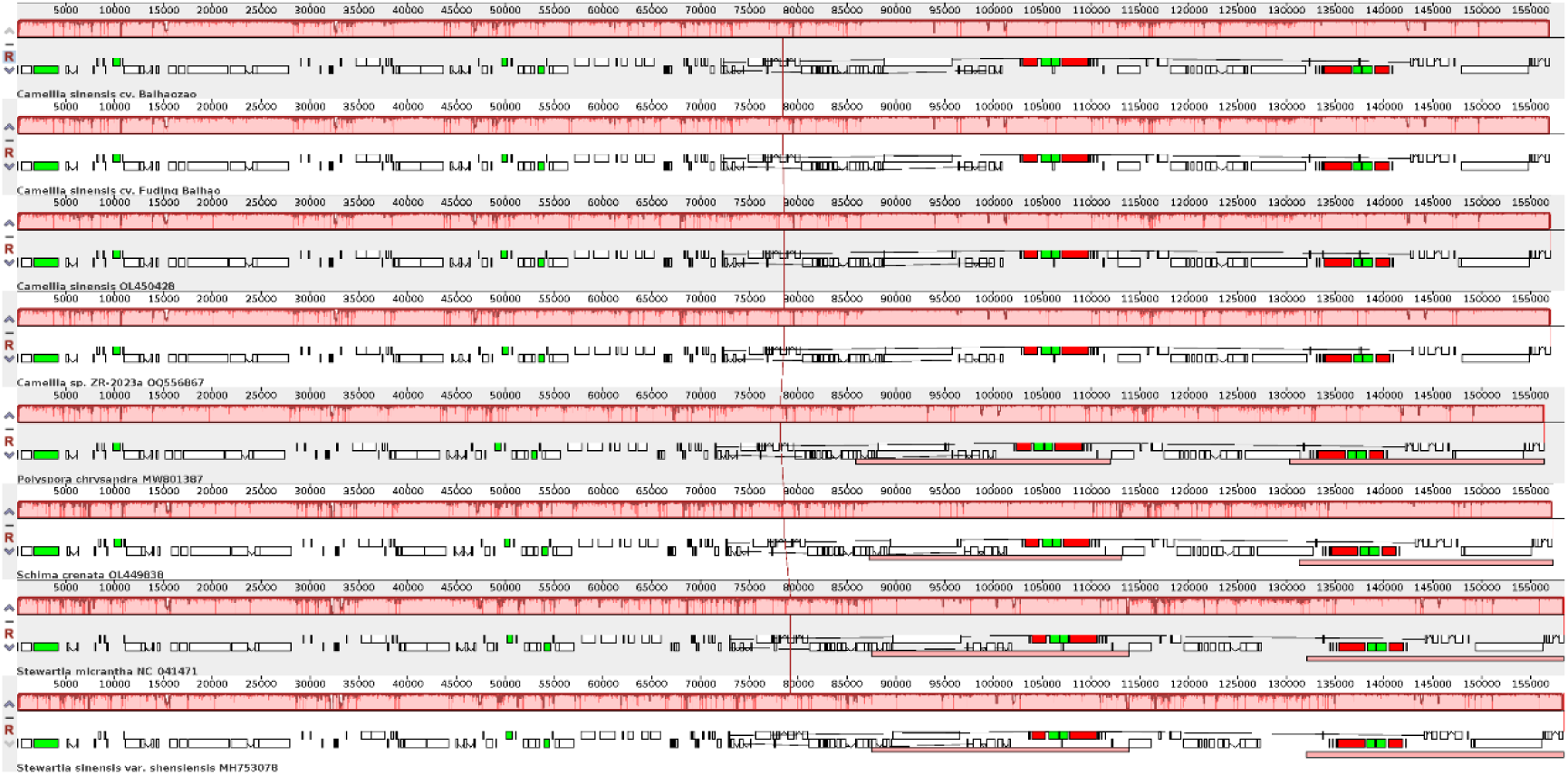
Homology analysis of cp sequences. Note: The short squares denote gene positions within the genome, with white squares representing coding sequences (CDS), green squares representing transfer RNA (tRNA), and red squares representing ribosomal RNA (rRNA). Colored blocks, referred to as Locally Collinear Blocks (LCBs), signify regions that are homologous to segments of another genome in the alignment. When an LCB is positioned above the centerline, the aligned region is in the forward orientation relative to the first genome sequence. Conversely, an LCB below the centerline indicates reverse complementarity alignment. Regions outside these blocks suggest a lack of homology between the input genomes.

### 2.7 Phylogenetic Inference

The cp genome plays a crucial role in phylogenetic studies. To elucidate the phylogenetic position of Baihaozao within the Camelliaceae family, a phylogenetic tree was constructed using cp sequences from 20 Camelliaceae species, with *Paeonia delavayi* (accession number KY817591) serving as the outgroup (Fig. 10). The resulting phylogenetic tree comprised 18 nodes, of which 13 exhibited 100% bootstrap support, thereby indicating a high level of reliability in the clustering outcomes. Within this phylogenetic framework, we assessed the phylogenetic affinities of Baihaozao in relation to the 20 species of *Camellia sinensis*. Theaceae can be taxonomically divided into the genera *Camellia*, *Polyspora*, *Stewartia*, and *Schima*. Notably, *Camellia* and *Polyspora* are sister taxa with high credibility. Within this framework, the relationship between Baihaozao and *Camellia sinensis* OL450428.1 is strongly supported, with both species residing on the same branch with 100% bootstrap support. This sub-branch is further nested within the same clade as *Camellia subintegra* NC_067087.1. Baihaozao exhibits a relatively distant phylogenetic relationship to the other three species within the genus *Camellia*. These findings suggest that Baihaozao is highly homologous to *Camellia sinensis* OL450428.1, while being more distantly related to other members of the genus.

**Fig. 10.**
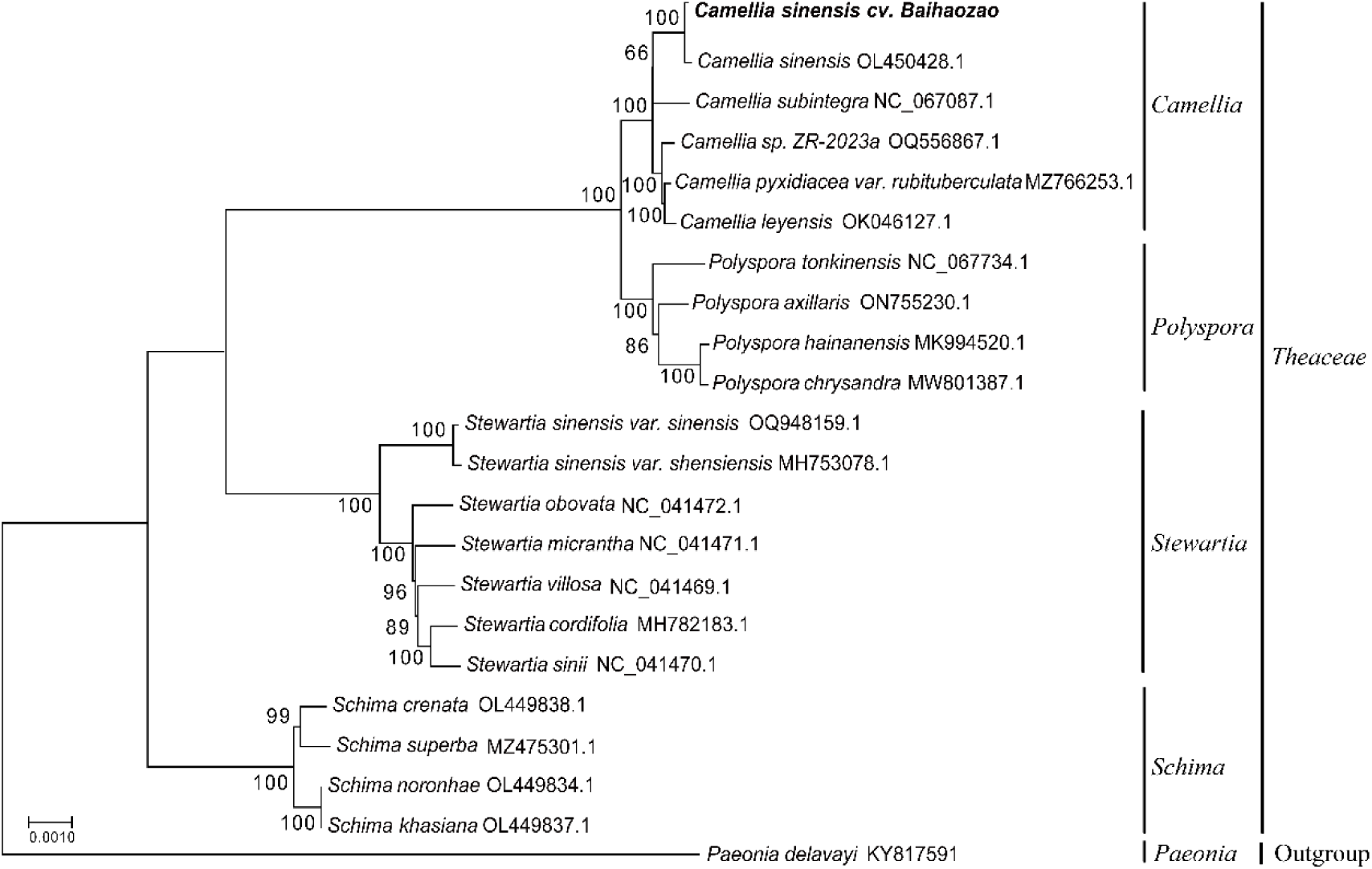
Phylogenetic analysis of cps. Note: The sequences are denoted by the Latin names of the respective species. The self-spreading value, which indicates the branching confidence of the evolutionary tree, is typically expressed as a number ranging from 0 to 100. The evolutionary branch length, also referred to as genetic variability or evolutionary distance, quantifies the extent of evolutionary divergence. A shorter branch length signifies a smaller degree of difference and a closer evolutionary relationship. This measure is usually represented as a decimal number between 0 and 1, corresponding to the magnitude of variation at a 100-base locus. (When the degree of genetic variation is almost zero, this data is not shown in this figure.)

## 3. Discussion

Cps, as fundamental organelles within plant cells, are indispensable for photosynthesis and various metabolic pathways[32]. These organelles are actively engaged in the synthesis of organic compounds and energy conversion processes, and they also harbor independent genetic material. In comparison to nuclear and mitochondrial genomes, the cp genome is characterized by several unique features: it possesses a relatively small genome size, a high gene copy number, highly conserved sequences, predominantly matrilineal inheritance, and haploid characteristics[33]. Given these unique characteristics, cp genomes have emerged as a critical research instrument for exploring plant genetic evolution, species classification and identification, DNA barcoding technology, phylogenetic analysis, and the phylogenetic reconstruction of plants[34, 35].

The cp genome sequences of over 130 species of *Camellia sinensis* (Camellia spp.) are presently cataloged in the National Center for Biotechnology Information (NCBI) database. Notably, the lengths of these cp DNAs exhibit minor variations, ranging from 150 to 160 kb, while consistently preserving a conserved quadripartite structure (SSC, LSC, IRa, IRb). This structural consistency is also observed in the cp genome of *Camellia* spp. Baihaozao, as reported in this study. In parallel, our investigation revealed that, akin to the majority of land plant cp genomes, the cp genome of Baihaozao demonstrates a highly conserved structural organization and gene composition. The cp genome spans 157,052 bp, closely resembling the cp genome size of the *Camellia sinensis* cultivar Zhuyeqi as reported in prior research[36]. Furthermore, a positive correlation was observed between GC content and sequence conservation, suggesting that regions with elevated GC content exhibit higher levels of conservation. The overall GC content of the Baihaozao genome was determined to be 37.30%, whereas the IR region exhibited a GC content of 42.95%. This disparity suggests that the IR region possesses greater structural stability compared to the LSC and SSC regions.

It is widely acknowledged that the contraction and expansion dynamics at the IR boundary influence the length of the cp genome. To elucidate these gene contraction and expansion phenomena, this study conducted a comparative analysis of the location and length of genes proximal to the IR boundary, specifically focusing on *rps19*, *rpl2*, *trnN*, *ndhF*, *ycf1*, and *trnH*. The results indicated that several genes were relocated at the IR boundary, with particular emphasis on the *ycf1* gene situated at the IR/SSC region boundary. This gene not only exhibited displacement but also demonstrated significant length variations. These alterations frequently precipitated the relocation of other genes at the IR boundary, culminating in genetic differences. These alterations frequently resulted in the displacement of other genes at the IR boundary, leading to genetic variations, which have been attributed to the presence of duplicated sequences within the *ycf1* gene[37]. The variations at the IR/LSC zone boundaries are minimal, with the exception of *Stewartia micrantha* NC 041471, which exhibits a more significant difference. This study indicates that the IR/LSC zone boundaries are more conserved compared to the IR/SSC zone boundaries. Furthermore, the contraction and expansion of the LSC/IR and SSC/IR zones not only influence their respective lengths but also serve as critical factors in the evolutionary development of cp[38]. The variations in the contraction and expansion of the IR zone boundaries across different species are of considerable significance for the study of cp IR zone boundaries. These variations can offer valuable insights for phylogenetic analysis and species identification[2].

SSRs are segments of the cp genome characterized by lengths of one to six nucleotide units[39]. These sequences are extensively utilized in plant species identification, genetic map construction, population evolutionary studies, and genetic diversity research of germplasm resources, owing to their notable attributes such as high abundance and polymorphism[40]. The study’s findings revealed the presence of five distinct nucleotide repeat units within the cp genome of Baihaozao, with single nucleotide repeat sequences being the most prevalent. It is noteworthy that these single nucleotide repeat sequences encompass not only A/T types but also G/C types. However, the occurrence of A/T type repetitive sequences (141) was significantly higher than that of G/C type sequences (16). This observation suggests that the elevated proportion of A/T base repeats may contribute to the high AT base content in the cp genome. Additionally, this study successfully identified several SSR loci, which can be utilized for genetic diversity analysis and phylogenetic studies in *Camellia sinensis*. Furthermore, dispersed repeat sequences, which constitute a significant component of the cp genome, harbor substantial genetic information[41]. The presence of these dispersed repeat sequences can result in DNA strand mismatches, thereby contributing to sequence variation[42]. Generally, organisms exhibiting higher evolutionary complexity tend to possess a greater abundance of dispersed repeat sequences[43]. In the analysis of dispersed repeat sequences within the cp genome of Baihaozao, an equal number of forward and palindromic repeats were identified, whereas no reverse or complementary repeats were detected. These dispersed repeat sequences exhibit potential utility as molecular markers in plant genetics research[44].

Codon preference is crucial in the decoding of genetic information, with specific usage patterns offering significant insights into gene function and species evolution[45, 46]. The cp proteome of Baihaozao comprises 26,471 genetic codons, with 64 codons employed to encode 20 amino acids. Methionine (Met) exhibited the highest codon usage, whereas tryptophan (Trp) had the lowest, suggesting a higher expression level of Met-related proteins in this Camellia species. In the interim, the codon AUG, commonly associated with encoding Met, demonstrated the highest RSCU frequency at 6.9566, underscoring its pivotal role in protein biosynthesis. This was closely followed by the UUA codon, which encodes leucine (Leu), exhibiting an RSCU value of 1.8984, indicating a substantial usage preference as well. In Baihaozao, the majority of codons exhibiting RSCU values exceeding 1.00 terminate in A or U bases. This pattern mirrors the codon usage preferences observed in the cp genomes of other tea plants, indicating a bias towards codons that are translated with higher accuracy and efficiency[47]. A comprehensive investigation into codon preferences across various organisms has the potential to enhance protein expression efficiency through optimized codon selection[48].

To investigate the phylogenetic affinities between Baihaozao and other genera within the Camellia family, we analyzed a total of 22 samples representing 21 species across four genera, along with one outgroup from the Camelliaceae family. This analysis was based on cp genome sequences (LSC+SSC+IR) and coding DNA sequences (CDS). Phylogenetic relationships within the Camelliaceae were elucidated using the Maximum Likelihood Evolutionary Tree (MLET) method. The findings indicated that within the four genera of Theaceae, Baihaozao was classified under the genus Camellia and exhibited the closest genetic relationship to *Camellia sinensis* OL450428.1. Furthermore, the genus Camellia demonstrated a closer phylogenetic relationship to *Polyspora*, while being more distantly related to *Stewartia* and *Schima*. The phylogenetic proximity between *Camellia* and *Polyspora* was notably closer compared to the more distant relationship observed between *Stewartia* and *Schima*. Additionally, the majority of branches in the molecular phylogenetic tree of *Camellia sinensis* received high support, underscoring the significant utility of the complete cp genome in resolving the phylogeny of complex taxa.

## 4. Conclusions

For the first time, the complete cp genome of *Camellia sinensis* sp. Baihaozao has been sequenced and characterized. The Baihaozao cp genome exhibits a typical tetrameric structure, encompassing a total length of 157,052 bp. The genome contains 133 predicted functional genes and has a GC content of 37.30%. Additionally, 247 SSRs were identified, with SNRs constituting the majority. A total of 247 SSRs were identified in the cp genome of Baihaozao, including 157 SNSRs, which were predominantly composed of A and T bases. Additionally, 32 codons in the Baihaozao cp genome exhibited a RSCU value greater than 1, with Leu being the most frequently utilized amino acid. Phylogenetic analysis indicates that Baihaozao is a distinctly monophyletic taxon, closely related to *Camellia sinensis* (GenBank accession number OL450428.1). These findings not only provide a scientific foundation for the development and utilization of Baihaozao camellia resources but also offer theoretical support for its conservation efforts.

## Abbreviations

cp: Chloroplast bp Base pair
LSC: Large single copy
SSC: Small single copy
SSR: Single sequence repeat IR Inverted repeat
CUB: Codon usage bias
RSCU: Relative synonymous codon usage
Pi: Nucleotide diversity
PCR: Polymerase chain reaction
Ka: Nonsynonymous locus
Ks: Synonymous locus
CDS: Cp-coding sequences
rRNA: Ribosomal RNA
tRNA: Transfer RNA
DNA: Deoxyribonucleic acid

## The contributions

**Zhiyin Chen:** Investigation, Data Curation, Methodology, Software, Validation, Writing - Original Draft, Funding Acquisition. **Youpeng Zhu:** Investigation, Data Curation, Software, Methodology, Validation. **Zhiming He:** Investigation, Data Curation, Software, Methodology, Validation. **Hongyu Li:** Investigation, Data. **Jing Huang:** Writing - Review & Editing. **Yihui Gong:** Conceptualization, Investigation, Data Curation, Methodology, Writing - Review & Editing, Visualization, Validation, Project Administration, Supervision, Resources.

## Data availability

The datasets generated and analyzed during this study are accessible in NCBI’s GenBank. Specifically, the complete cp genome sequence of Baihaozao has been deposited under accession number PQ066596. The accession numbers for the additional datasets utilized and analyzed in this study are detailed in the “Methods” section.

## Declarations

### Ethics Approval and Consent to Participate

No specific license was required for the collection of samples in this study. The study was conducted in compliance with relevant Chinese legislation.

